# Marine deep biosphere microbial communities assemble in near-surface sediments in Aarhus Bay

**DOI:** 10.1101/557041

**Authors:** Caitlin Petro, Birthe Zäncker, Piotr Starnawski, Lara M. Jochum, Timothy G. Ferdelman, Bo Barker Jørgensen, Hans Røy, Kasper U. Kjeldsen, Andreas Schramm

## Abstract

Analyses of microbial diversity in marine sediments have identified a core set of taxa unique to the marine deep biosphere. Previous studies have suggested that these specialized communities are shaped by processes in the surface seabed, in particular that their assembly is associated with the transition from the bioturbated upper zone to the nonbioturbated zone below. To test this hypothesis, we performed a fine-scale analysis of the distribution and activity of microbial populations within the upper 50 cm of sediment from Aarhus Bay (Denmark). Sequencing and qPCR were combined to determine the depth distributions of bacterial and archaeal taxa (16S rRNA genes) and sulfate-reducing microorganisms (*dsrB* gene). Mapping of radionuclides throughout the sediment revealed a region of intense bioturbation at 0-6 cm depth. The transition from bioturbated sediment to the subsurface below (7 cm depth) was marked by a shift from dominant surface populations to common deep biosphere taxa (e.g. Chloroflexi & Atribacteria). Changes in community composition occurred in parallel to drops in microbial activity and abundance caused by reduced energy availability below the mixed sediment surface. These results offer direct evidence for the hypothesis that deep subsurface microbial communities present in Aarhus Bay mainly assemble already centimeters below the sediment surface, below the bioturbation zone.

## 1. Introduction

Marine sediments are a massive microbial habitat, harboring as many as 5.39 × 10^29^ prokaryotic cells on a global scale and 0.18-3.6% of Earth’s total living biomass (Kallmeyer et al., 2012; Parkes et al., 2014). Microorganisms deposited on the sediment surface are gradually buried into the seabed as sedimenting particulate matter accumulates on the seafloor. This burial can isolate microorganisms from the surface world for hundreds to millions of years, cutting populations off from fresh detrital organic matter and subjecting them to severe energetic limitations that increase with sediment depth (Langerhuus et al., 2012; Lomstein et al., 2012; Middelburg, 1989). Despite these energy limitations, deep marine sediments are populated by living microbial communities that persist down to 2.5 km below the seafloor (reviewed in Parkes *et al*., 2014; Inagaki *et al*., 2015). Phylogenetic diversity (16S rRNA gene) analyses of the subsurface environment have identified a core set of uncultivated microbial taxa that have become synonymous with the marine deep biosphere (Parkes *et al*., 2014; Carr *et al*., 2015; Orcutt *et al*., 2011; Kubo *et al*., 2012; Biddle *et al*., 2006; Teske, 2006). But at which sediment depth and by which processes these deep subsurface communities are formed remains unclear.

Work thus far has suggested that selective survival is the major driver of microbial community assembly in subsurface marine sediments (Jochum et al., 2017; Petro et al., 2017; Starnawski et al., 2017; Walsh et al., 2015). Selection filters out populations during burial, creating a deep subsurface biosphere that is populated by rare descendants of the surface sediment community (Jochum et al., 2017; Starnawski et al., 2017). Persisting taxa exhibit little genetic diversification across sediment depths, suggesting that adaption to deep biosphere conditions plays a limited role in driving the assembly of the deep subsurface community (Starnawski et al., 2017). The lack of adaptive evolution is likely due to the exceedingly long generation times encountered within marine sediments, and thus the limited number of generations, which limit the potential for genetic change during burial (reviewed in Jørgensen and Marshall, 2016; Lever *et al*., 2015a).

Shortest generation times have been estimated near the sediment surface, owing to the availability of freshly deposited organic matter and high potential electron acceptors (e.g. oxygen and nitrate), which collectively promote community growth and turnover (Hoehler and Jørgensen, 2013; Langerhuus et al., 2012; Lomstein et al., 2012). Microbial activity is further stimulated near the sediment surface due to bioturbation by benthic macrofauna. Bioturbation processes include faunal burrow construction and maintenance, both of which continually rework and irrigate the upper 10 ± 5 cm of the seabed (Boudreau, 1998; Meysman et al., 2006). While bioturbation occurs globally, both the intensity of bioturbation and the depth of mixing are influenced by factors such as seasonality and depth of the overlying water column (Teal et al., 2008). Water depth was shown to play a minor role in driving differences in bioturbation between sites, indicating that the intensity and depth of mixing are only marginally different between coastal and deep ocean sediments. In addition to enhancing the dispersal of microbial cells, bioturbation also increases microbial energy availability by introducing labile organic matter and oxygen from the overlying water column into the seabed (Kristensen et al., 2012; Kristensen and Holmer, 2001). Bioturbation has been shown to influence microbial communities near the seafloor, most notably by driving the dominance of Bacteria relative to Archaea (Chen et al., 2017) that has been observed repeatedly in surface sediments (Giovannelli et al., 2013; Lipp et al., 2008).

Below the bioturbation zone, sediments exhibit a vertical geochemical zonation that results from the thermodynamically-driven succession of available electron acceptors in the sediment (Canfield et al., 1993; Froehlich et al., 1979). In organic-rich coastal marine sediments, the sulfate reduction (SR) zone occurs immediately below the bioturbation zone (Jørgensen, 1982). Here, sulfate is the main electron acceptor for microbial respiration. Below the SR zone, the sulfate-dependent anaerobic oxidation of methane takes place within a narrow zone defined as the sulfate-methane transition (SMT) (Knittel and Boetius, 2009; Leloup et al., 2007; Thomsen et al., 2001). Each biogeochemical process is mediated by a unique guild of terminal oxidizers, such as sulfate-reducing microorganisms (SRM) and methanogens, respectively.

Analyses of subsurface microbial diversity across geochemical zones in Aarhus Bay sediments have shown that communities change most dramatically between the surface environment and the SR zone below (Chen et al., 2017; Jochum et al., 2017; Starnawski et al., 2017). Below the SR zone, a large fraction of the microbial community is more stable and persists with burial into the deep subsurface. It was therefore suggested that the assembly of subsurface communities is associated with the transition from the bioturbated surface zone of the sediment to the unmixed sediment zones below (Jochum *et al*. 2017; Starnawski *et al*. 2017). To test this hypothesis, we performed a fine-scale analysis of the distribution and activity of microbial populations present within the upper 50 cm of sediment from Aarhus Bay (Denmark). Because patterns of microbial diversity across sediment depths in Aarhus Bay are highly reproducible between sampling sites (Chen *et al*., 2017; Starnawski *et al*., 2017; Jochum *et al*., 2017), we chose to explore these patterns at high-resolution within sediment collected from one well-studied sampling site, station M5 (Langerhuus et al. 2012; Chen *et al*. 2017). Sediment cores were approached as a vertical transect, with the aim of examining population dynamics associated with burial of resident microbial communities over time. Next-generation sequencing was employed to determine the distributions of bacterial and archaeal taxa (16S rRNA gene) as well as SRM, using the the gene encoding the beta subunit of the dissimilatory (bi)sulfite reductase (*dsrB*) as a marker gene (Müller et al., 2014). This approach was utilized in order to compare population dynamics of the total microbial community with a known guild of terminal oxidizers (SRM). Sequencing was coupled with quantitative PCR (qPCR) and total cell counts in order to map changes in the absolute abundances of dominant taxa at each depth. Sulfate reduction rates were measured as a proxy for microbial activity and overall organic carbon mineralization in the sediment and the vertical extent of the bioturbation zone was mapped by determining radionuclide distributions.

## 2. Experimental Procedures

### Sampling

A sediment core was taken by Rumohr Corer during a cruise with the *RV Aurora* in September 2014 at Aarhus Bay (Denmark) station M5 (56° 06.333′N, 10° 27.800′E; water depth 27 m). This core was used for all subsequent analyses unless otherwise noted. The sediment core was stored and sampled at 15°C, corresponding to the *in situ* temperature of the bottom water. Subsamples for DNA extractions, sulfate reduction rate measurements, and cell counts were collected with sterile, pre-cut 2.5 ml syringes from the intact core. Samples were taken in 1 cm increments from the top down to 40 cm, with three additional deeper reference samples at 45, 48 and 50 cm depth.

In order to assess the reproducibility of our results across multiple cores and sampling times, we took four additional Rumohr cores from site M5 in both 2017 and 2018. The processing and analysis of these cores is decribed in the Supplementary Material.

### Sulfate reduction rates (SRR)

SRR were determined in 1 cm intervals using the ^35^S-tracer technique (Røy et al. 2014). Subsamples of 2.0 cm^3^ were taken with cut-off syringes that had been covered with Parafilm® and inserted cut-side down into anoxic marine sediment to prevent the samples from becoming oxygenated during processing. 0.5 cm^3^ of sediment from each subsample was transferred into a microcentrifuge tube and pelleted by centrifugation. The supernatant was collected and used to measure the sulfate concentration in the pore water by Ion Chromatography (Dionex IC 2000, Thermo Scientific, Dreieich, Germany). The remaining sediment was injected with 10 µl of ^35^S-SO_4_^2-^ radiotracer (20 kBq µl^−1^) and incubated for 5 h in the dark at 15°C under anoxic conditions. To stop the incubations, the samples were transferred into 5 ml of 20 % Zn-acetate solution and stored at −20°C. Total reduced inorganic sulfur was separated from sulfate and reduced to sulfide by the cold chromium distillation procedure (Kallmeyer *et al*., 2004 with modifications by Røy *et al*. 2014). The radioactivity was measured on a Packard Tri-Carb 2900 TR Liquid Scintillation Analyzer (PerkinElmer, Waltham, MA). Blanks consisting of sediment killed with 20 % Zn-acetate prior to radiotracer injection were measured in parallel. The background radioactivity of the equipment and reagents was tested using samples containing only 5 ml Zn-acetate and Ecoscint XR (BioNordika, Herlev, Denmark). SRR were calculated according to Jørgensen (1978).

### Distribution of ^210^Pb (excess) and ^137^Cs

Sediment samples for determination of ^210^Pb, ^226^Ra, ^137^Cs, and ^40^K distributions were derived from a separate, 51 cm long core collected from station M5 in April 2016. The core was extruded from the liner and cut into 1 cm sections. The sections were dried at 105°C for 2 days, finely ground, and precisely weighed into polysulfone screw-top containers with approximately 20 g of sediment per sample. The samples were sealed with electrical tape and stored for 20 days to allow ^222^Rn ingrowth to secular equilibrium before counting. The radioactivities of ^210^Pb (46.5 keV), ^214^Pb (295 and 352 keV), ^214^Bi (609 keV), ^137^Cs (662 keV) and ^40^K (1405 keV) were then measured by low-level gamma-spectroscopy (Eurysis Ge Coaxial Type N detector, Canberra Industries, Rüsselsheim, Germany). A detector efficiency-energy curve was calibrated using a certified reference U-Th ore (Canmet DL-1a) and laboratory grade KCl (for ^40^K). Sediment self-absorption at 45.5 keV was corrected according to the method of Cutshall *et al*., (1983). The activity of ^210^Pb supported by the sediment’s natural contents of daughters of the ^238^U decay chain was estimated from the activity of ^226^Ra. ^226^Ra was estimated from daughter nuclide activities of ^214^Pb (295 and 352 keV), ^214^Bi (609 keV), assuming secular equilibrium between ^226^Ra, ^222^Rn, ^214^Bi and ^214^Pb. ^226^Ra- supported ^210^Pb was then subtracted from the total ^210^Pb activity to estimate the excess ^210^Pb derived via deposition.

### Calculation of bioturbation

A numerical model was set up to calculate the magnitude of sediment mixing by reverse modeling of the distributions of ^137^Cs and excess ^210^Pb in the upper 36 cm of sediment. A constant flux of ^210^Pb, corresponding to the depth-integrated rate of measured excess ^210^Pb activity, was imposed across the sediment surface. During runtime ^210^Pb was allowed to decay with a half-life of 22.3 years. Sedimentation was simulated by imposing downwards advection at 0.96 mm year^−1^ according to the mean sedimentation rate at station M5 (derived from ^14^C age determination of bivalve shells, Langerhuus *et al*., 2012). The model was divided into two depth-domains with independently controlled rates of bioturbation to simulate the distinct change in bioturbation between the upper 5.75 ± 5.67 cm “mixed zone” and the deeper sediment (Teal et al., 2008). The model was allowed to run until steady state. The division-depth between the upper and the lower zone, and the rate of mixing in the two zones, were then adjusted iteratively to give the best fit to the measured distribution of excess ^210^Pb activity.

A second iteration of the model was initiated with one unit of ^137^Cs in the upper one mm of the sediment, simulating the deposition of fall-out from the Chernobyl nuclear accident. ^137^Cs was then allowed to decay with a half-life of 30.17 years and allowed to move by the same bio-diffusion and convection as ^210^Pb. This iteration of the model was run for 29 years, corresponding to the time from the Chernobyl nuclear accident in April 1986 to the sediment core collection in April 2016. The rates of bioturbation in the two depth-zones were again adjusted iteratively to simultaneously achieve the best steady-state simulation of the measured depth-distributions of ^210^Pb and the best transient distribution of ^137^Cs activity. The model was implemented in Comsol Multiphysics.

### Total cell counts (TCC)

Sediment subsamples of 1 mL were preserved 1:2 (v/v) in 4 % Paraformaldehyde (w/v) and stored at 4°C until further processing. The fixed samples were homogenized and cells detached from sediment particles according to the protocol by Lavergne *et al*. (2014). The cells were stained with SYBR Gold (1 µl 10,000X in 10 ml 1X PBS) and 4,6-diamidino-2-phenylindole (DAPI) (1 µg ml^−1^) and analyzed by epifluorescence microscopy (Zeiss Axiovert 200, Zeiss, Jena, Germany).

### DNA extractions

Sediment subsamples for DNA extraction were stored at −80°C prior to processing. To avoid contamination from the sampling equipment, sediment that was in contact with the core liner was discarded. Approximately 0.2 g of sediment was used for each DNA extraction, which included an initial washing step to remove extracellular DNA (Lever et al., 2015b). After washing, cells were lyzed by enzymatic treatments (Kjeldsen et al., 2007) followed by incubation at 65°C for 2 hours in 2% sodium dodecyl sulfate. At this point, the DNA was extracted using the FastDNA Spin Kit for Soil (MP Biomedicals, Holte, Denmark) according to the manufacturer’s instructions.

### Quantification of 16S rRNA and dsrB genes

Bacterial and archaeal 16S rRNA genes were quantified by qPCR according to Starnawski *et al*. (2017) using the primer pairs Bac908F/Bac1075R (Ohkuma and Kudo, 1998) and Arch915Fmod/Arch1059R (Cadillo-Quiroz et al., 2006), respectively. SRM abundance was estimated by qPCR of the *dsrB* gene as described by Jochum *et al*. (2017). DNA extraction efficiency was calculated by comparing TCC to 16S rRNA gene copy numbers generated from qPCR, assuming that bacteria and archaea on average harbor 4.1 and 1.6 16S rRNA gene copies cell^−1^, respectively (Stoddard et al., 2015). Using this approach, we estimated an average DNA extraction effiency of 7% for all depths.

### Ion Torrent PGM sequencing and analysis

The bacterial and archaeal 16S rRNA genes were amplified using the primer pair Univ519F/Univ802R (Wang and Qian, 2009). The resulting ∼283-bp long fragment was barcoded and sequenced on an Ion Torrent PGM using 300 bp chemistry (Life Technologies, Carlsbad, CA). Amplification, sequencing, and sequence analysis of the 16S rRNA gene were all done according to Starnawski *et al*. (2017). Reads were clustered into operational taxonomic units (OTUs) with a 97% similarity cutoff using the UPARSE-OTU algorithm implemented in the usearch v7.0.959_i86osx32 software (Edgar, 2013). Representative sequences of each of the 2431 OTUs identified by 16S rRNA gene sequencing were deposited in GenBank under accession numbers MG637451 - MG639881.

*DsrB* sequencing was performed according to Jochum *et al*. (2017) using the primer variant mixtures dsrB-F1a-h and dsrB-4RSI1a-f (Lever, 2013). The resultant *dsrB* sequence libraries were filtered and analyzed using the pipeline described in Jochum *et al*. (2017). Species-level OTUs were clustered at a 90% similarity cutoff using pick_otus.py from the QIIME package (Caporaso et al., 2010). The taxonomic identity of SRM was resolved by classifying translations of quality-filtered reads according to the taxonomic framework proposed by Müller *et al*. (2014), with modifications and procedures described by Jochum *et al*. (2017). Representative OTU sequences were deposited in GenBank under the accession numbers MG742726 – MG744217.

The classified OTUs were transferred to the R statistical environment (Version 3.4.2), where all additional analyses were performed. Non-metric multidimensional scaling (NMDS) ordinations were calculated on the OTU level, both for the total microbial community and the SRM community. Prior to ordination, the OTU abundance tables were randomly subsampled to even sequencing depth using the rarefy_even_depth function in the phyloseq package (McMurdie and Holmes, 2013). The NMDS coordinates were calculated from the subsampled datasets using the metaMDS function implemented in the vegan package (Oksanen et al., 2017).

### Estimates of biomass turnover times

SRR measurements were combined with qPCR data to estimate per-cell metabolic rates (csSRR) and microbial biomass turnover times. For these calculations, *dsrB* gene copy numbers were used to estimate the abundance of SRM, assuming one *dsrB* gene per genome and a DNA extraction efficiency of 7% (Wagner et al., 2005). Biomass turnover times (T_b_) were estimated by dividing the biomass present within a given depth by the rate of biomass production. The biomass was calculated by multiplying the number of SRM (cells per g sediment) with the amount of carbon contained within a single cell (assuming an average cellular carbon content of 21.5 fg C per cell as empirically determined for subseafloor cell populations (Braun et al., 2016)). The rate of production was calculated from the measured SRR (nmol g^−1^ sediment d^−1^) and an estimated growth yield (8% according to D’Hondt *et al*. (2014)), assuming 2 moles of organic carbon oxidized per mole of sulfate reduced. The cumulative generation times per depth interval were calculated using an estimated sedimentation rate of 0.96 mm year^−1^ (Langerhuus et al., 2012).

## 3. Results

### Geochemistry and microbial activity

Pore water sulfate concentrations at site M5 decreased steeply with sediment depth from ∼27 mM at 3 cm below seafloor (cmbsf) to ∼2 mM at the deepest analyzed coring depth of 50 cmbsf (Figure 1a). Alignment of the sulfate concentrations with a methane profile taken previously from the same site revealed a sulfate methane transition (SMT) between 45 and 55 cmbsf. The sulfate and methane profiles were reproduced in the cores taken in 2017 and 2018, indicating that the geochemical conditions at site M5 are stable over time (Figure S6). Maximum rates of sulfate reduction (177 - 218 nmol cm^- 3^ d^−1^) occurred at 1-3 cmbsf (Figure 1b). Below 3 cmbsf, the SRR dropped steeply with depth, reaching ∼ 0.4 nmol cm^-3^ d^−1^ at 50 cmbsf. Much like the sulfate and methane profiles, this pattern was highly reproducible across years (Figure S6).

**Figure 1.**
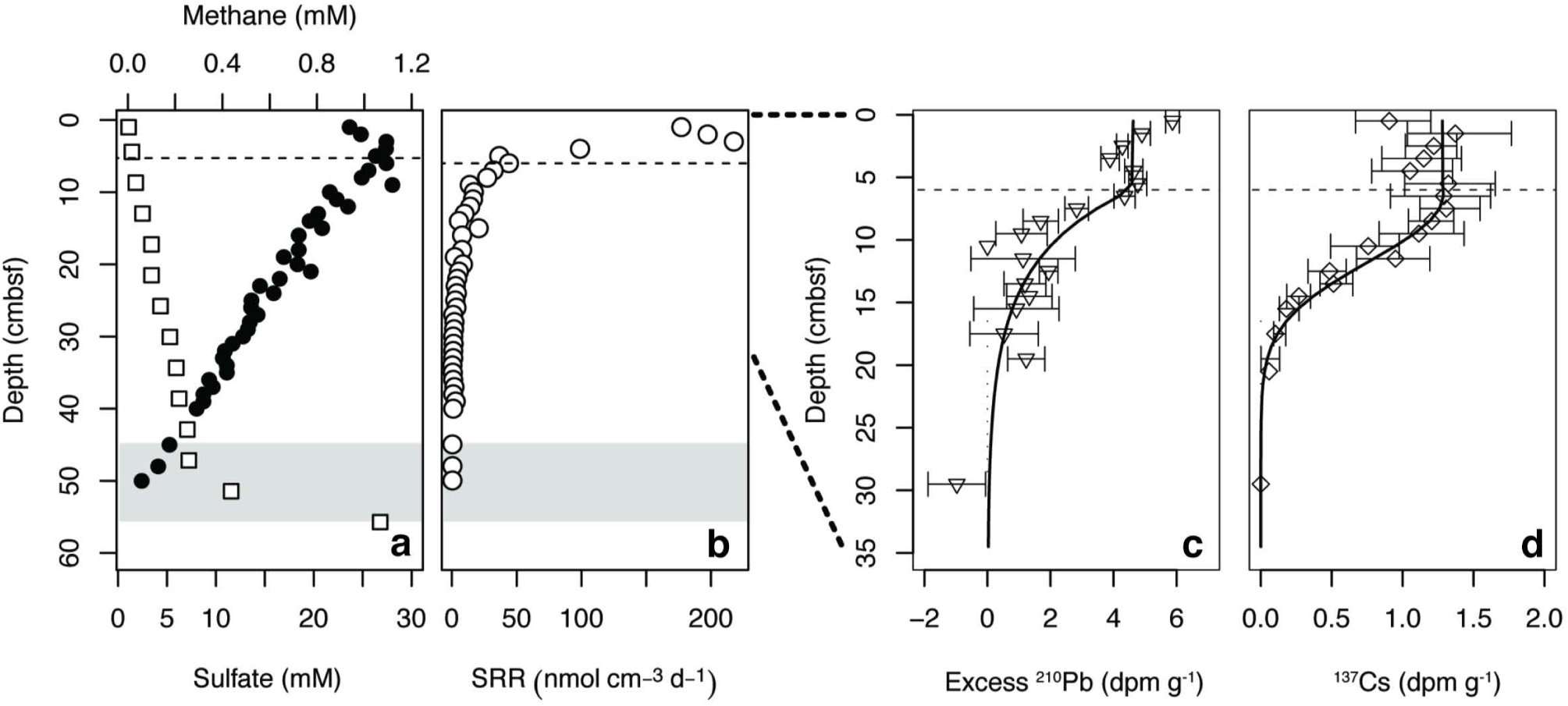
(a) Geochemical zonation of site M5, Aarhus Bay (Denmark). The sulfate profile (•) was measured in the present study, while the methane profile (□) was obtained from previous work at the same site (Hans Røy, unpublished). (b) Sulfate reduction rates (SRR). (c) Profile of excess ^210^Pb and (d) ^137^Cs in the surface of site M5. Error bars represent 1 sigma standard deviation of gamma counting uncertainty. Gray shading indicates the onset of the sulfate methane transition (SMT). The dashed lines indicate the bottom of the highly mixed surface layer, or bioturbation zone, based on modeling of excess ^210^Pb and ^137^Cs.

### Sediment mixing

According to measurements of excess ^210^Pb and ^137^Cs activity a region of intense mixing is present from 0-6 cmbsf (Figure 1). Below this depth range, ^210^Pb activity dropped logarithmically, but penetrated deeper than what would be expected from the balance between burial and decay. The effects of mixing were even more evident with ^137^Cs originating from the Chernobyl nuclear accident, which should be found as a distinct peak at 2.3 cm below the sediment surface in the absence of sediment mixing (Moros et al., 2017). Mixing in the upper 6 cm alone would increase penetration to maximum 8.3 cm, but the radioisotope was present at least to 20 cmbsf. Reverse modeling of the depth distribution of ^210^Pb and ^137^Cs suggested a bio-diffusion coefficient of at least 10^-2^ m^2^ year^−1^ in the upper 6 cm, and near 4 x10^-5^ m^2^ year^−1^ in the zone between 6 and 35 cmbsf.

### Vertical distribution of Bacteria and Archaea

TCC and qPCR data showed a sharp drop in microbial abundance between the sediment surface and the bottom of the zone of more intense bioturbation (0-6 cmbsf) referred to as the “bioturbation zone” below (Figure 2). Within this region, TCC dropped by 65%, while bacterial and archaeal 16S rRNA gene copy numbers dropped by 86% and 70%, respectively. 16S rRNA gene copy numbers followed the same distribution exhibited by TCC, but with lower abundances (Figure 2).

**Figure 2.**
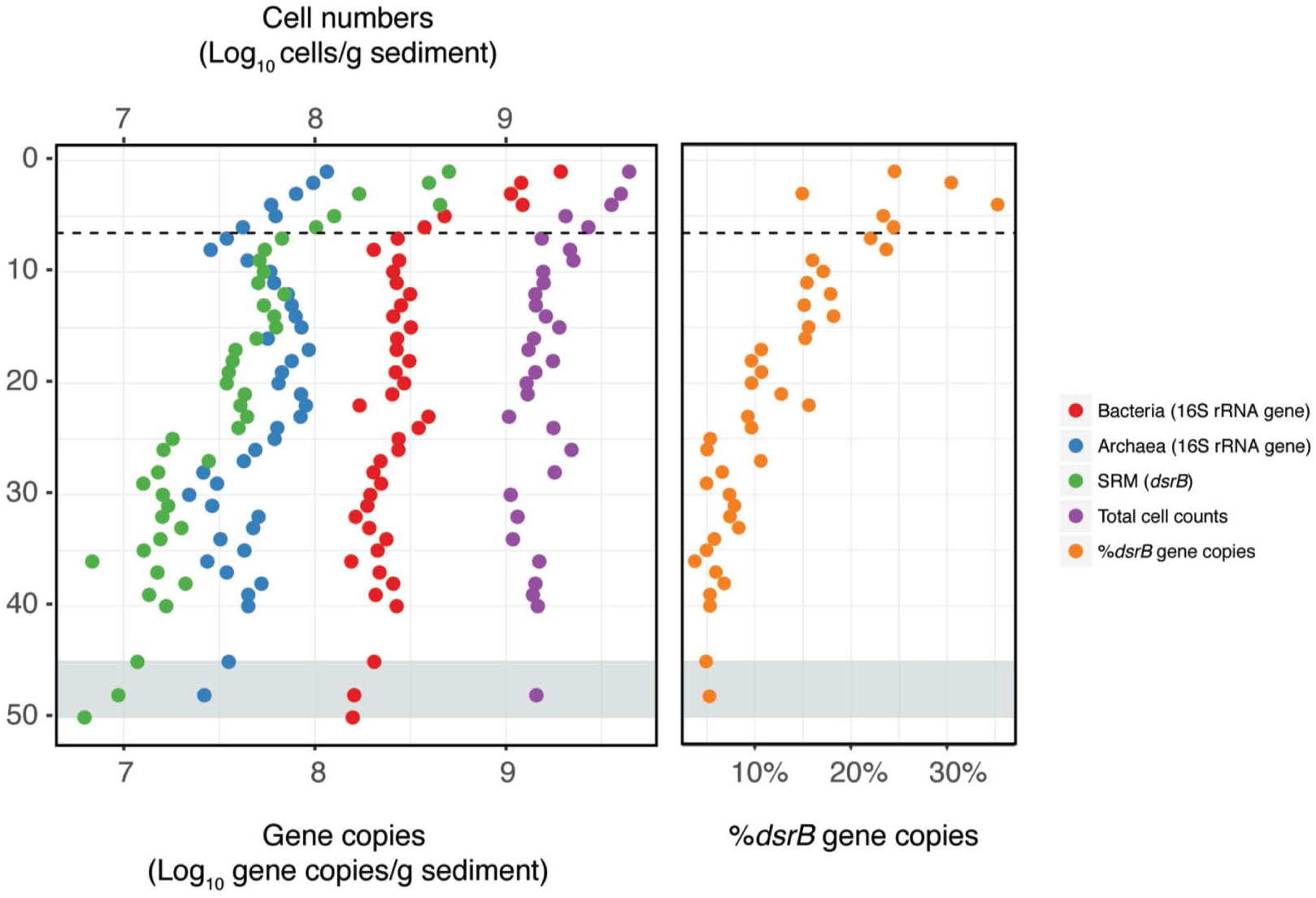
Distribution of microbial abundances with depth. Bacterial, archaeal, and sulfate reducing microorganisms (SRM) were quantified by qPCR. Total cell counts were quantified by direct epifluorescence microscopy of microbial cells. The dashed line indicates the depth of the bottom of the bioturbation zone and the gray shading indicates the onset of the sulfate methane transition (SMT).

Non-metric multidimensional scaling (NMDS) ordination of 16S rRNA gene sequence OTU distributions revealed that the most pronounced change in OTU composition occurred between 3 and 7 cmbsf, corresponding exactly to the bottom of the bioturbation zone (Figure 3a). Below the bioturbation zone samples clustered together, indicating that they harbored more similar communities when compared to the communities present within the bioturbation zone. Samples assigned to the subsurface (SR, 7-36 cmbsf) and the bottom of the core (SMT, 48-50 cmbsf) also clustered together, suggesting that the composition of the microbial community below the bioturbation zone is relatively stable and changes only gradually with depth (Figure 3a).

**Figure 3.**
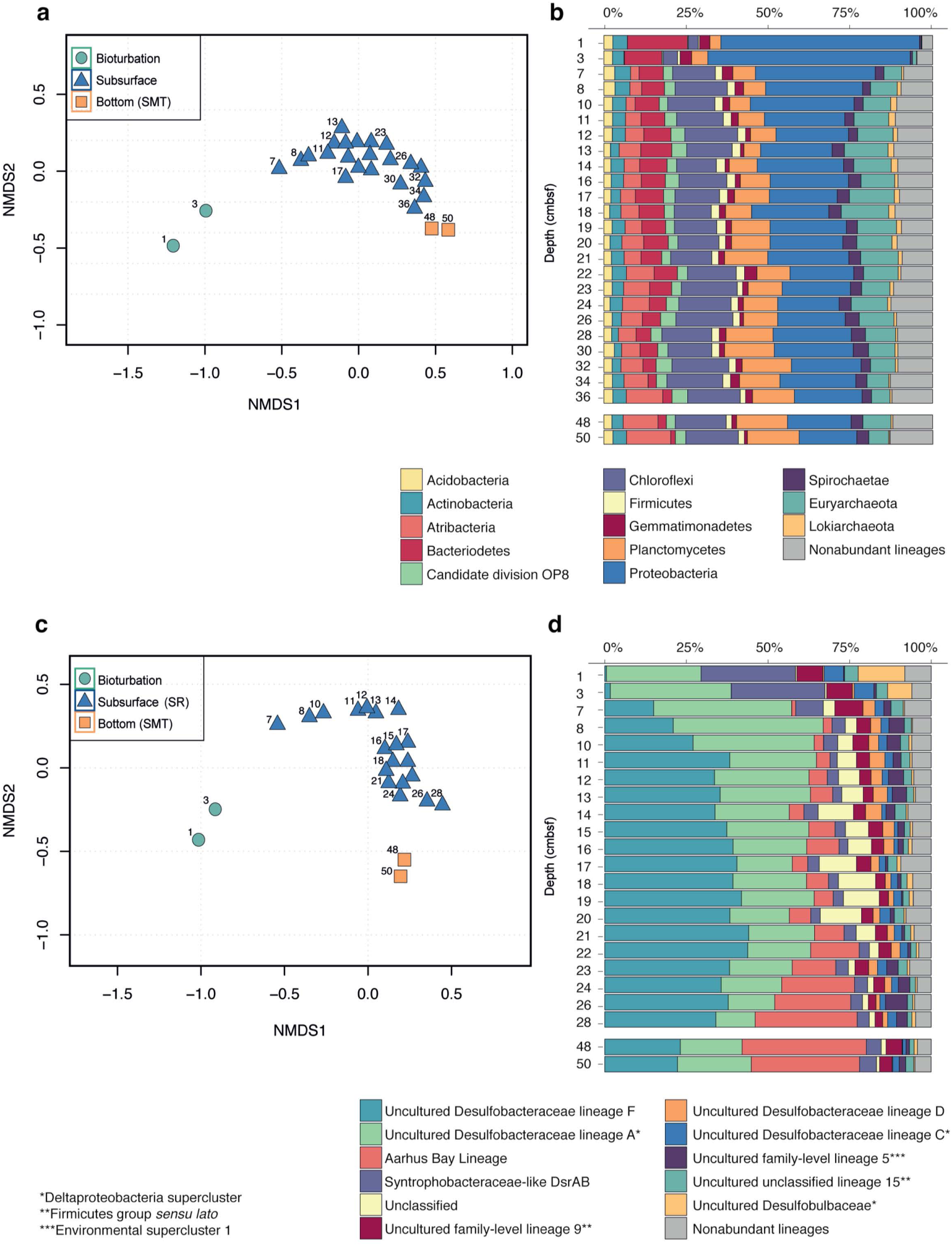
Changes in the microbial community with depth. (a & c) Non-metric multidimensional scaling (NMDS) of subsampled sequencing datasets for all Bacteria and Archaea (a) and SRM lineages (c). NMDS ordinations were calculated at the OTU level (b & d) Stacked bar plots of the relative sequence abundances for the total community (b) and the SRM lineages (d).

Proteobacteria sequences were negatively associated with depth (Spearman, r = −0.73, P < 0.001) and dropped in relative abundance from 60% in the bioturbation zone to 30% at 7-8 cmbsf and 18% at 50 cmbsf (Figure 3b). The drop in Proteobacteria sequences was primarily driven by sequences belonging to the Gammaproteobacteria, which dropped from 30% in the bioturbation zone to 1-10% in the subsurface below (Supplementary, Figure S1).

In contrast to the Proteobacteria, sequences clustering within the Chloroflexi were positively associated with depth (Spearman, r = 0.57, P < 0.01) and increased in relative abundance from 3% at 3 cmbsf to 16% at 8 cmbsf (Figure 3b). The increase in relative abundance of Chloroflexi was primarily driven by OTUs belonging to the class Dehalococcoidia, which increased from <1% of total sequencing reads in the surface to 12% at 50 cmbsf (Supplementary, Figure S1). A similar positive correlation with depth was observed for sequences clustering within the Atribacteria (Spearman, r = 0.84, P < 0.001), which comprised <0.01% of total sequences in the bioturbation zone. A small community of Atribacteria began to develop immediately below the bioturbation zone (3% of total sequences at 7 cmbsf), and increased to 13% at 50 cmbsf (Figure 3b).

Major taxonomic shifts seen for dominant phyla were also observable on the OTU level. Mapping of the 25 most abundant OTUs at different depths throughout the core revealed a dramatic change in the OTU community composition right below the bioturbation zone (7 cmbsf), from a community dominated by Proteobacteria to more diverse high-ranking OTUs belonging to the Chloroflexi, Atribacteria, and several archaeal phyla (Supplementary, Figure S2). The OTUs that appeared right below the bioturbation zone (7-10 cmbsf) continued to increase in rank with increasing sediment depth. By 50 cmbsf, the 25 most dominant OTUs comprised 45% of the total sequencing reads, with a single Atribacteria OTU accounting for 12% of the total microbial community. Similar taxonomic shifts within bacterial phyla were also seen in the cores taken from 2017, including a marked increase in the relative abundances of Atribacteria and Chloroflexi occurring between 5 and 10 cmbsf (Figure S7A). Our separate analyses of archaeal lineages from the cores taken in 2018 also show changes in the archaeal community across depths (Figure S7B). Most notably we see an increase in the relative abundance of Lokiarchaeia with depth, a class within the newly characterized Asgardaeota phylum (Spang et al., 2015; Zaremba-Niedzwiedzka et al., 2017). There is also a stark decrease in the relative abundance of Thaumarchaeaota, specifically the class Nitrososphaeria, with increasing depth.

In a separate analysis, we traced OTUs across depths in order to identify OTUs from the surface that persisted to the bottom of the core (Supplementary, Figure S3). This revealed 92 OTUs that were present in every sequenced depth, from the bioturbation zone to 50 cmbsf. While this set of persisting OTUs represented only 12% of the total OTU richness at 50 cmbsf, they comprised 45% of the total sequencing reads (Figure S3). By converting OTU relative abundances to absolute abundances using qPCR data (Starnawski et al., 2017), we found that several OTUs also increased in absolute abundance with depth, running counter to the drop in absolute abundance observed for the total microbial community and the majority of dominant surface OTUs (Figure 4). The OTUs which increased in absolute abundance with depth (Supplementary Table 1) belong to common subsurface lineages such as the Atribacteria and members of the class Phycisphaerae, both of which have been found to constitute a significant portion of the microbial community hundreds of meters below the seafloor (Petro et al., 2017).

**Figure 4.**
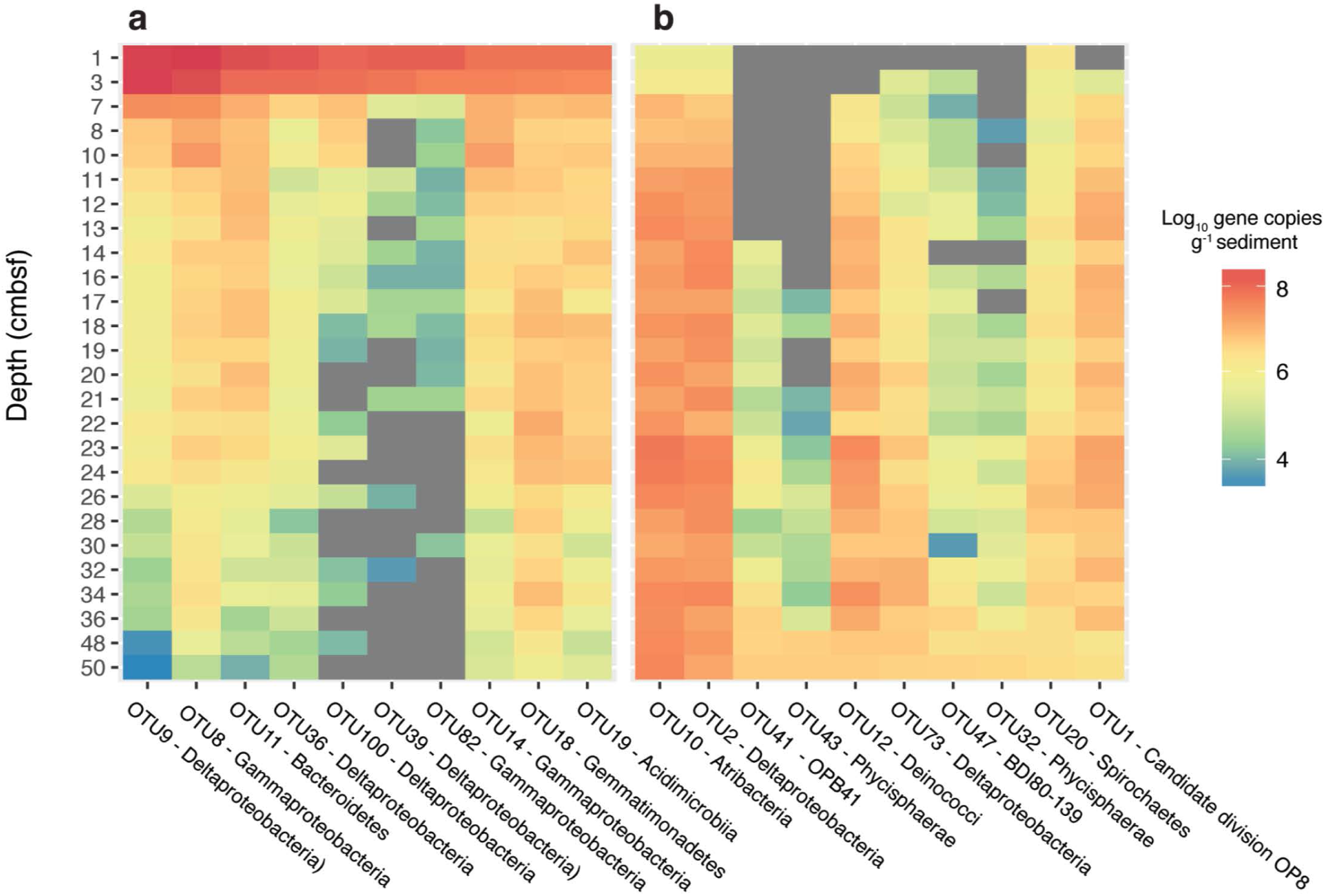
Estimated absolute abundances of the most dominant OTUs identified by sequencing of 16S rRNA gene sequencing. Absolute abundances were calculated by multiplying relative sequences abundances by gene copy numbers obtained from qPCR of the 16S rRNA gene, assuming an average of 4.1 and 1.6 16S rRNA operons per cell for bacteria and archaea, respectively. The top ten most abundant OTUs present at 3 cmbsf are displayed in (a) and the top ten most abundant OTUs present at 50 cmbsf are displayed in (b). The color represents the log_10_ absolute abundance of each OTU. Grey areas represent depths where no sequences were recovered for the specified OTU.

### Vertical distribution of sulfate reducing microorganisms (SRM)

Both qPCR and sequencing analysis of *dsrB* genes indicated that SRM were present throughout the entire depth profile (Figures 2 & 3d). *DsrB* gene copy numbers equated to 5-30% of the 16S rRNA gene copy numbers, with the highest relative abundance occurring within the bioturbation zone (Figure 2). The *dsrB* gene copy number showed a strong positive correlation to the sulfate reduction rate (R^2^ = 0.86, P < 0.001; Figure S4), highlighting *dsrB* gene quantification as a means to estimate SRM abundance. Furthermore, the depth-associated pattern of *dsrB* gene copy numbers could be reproduced in cores sampled in 2017 and 2018 (Figure S8).

Sequencing analysis of the *dsrB* gene grouped the OTUs into 44 different lineages. NMDS ordination of the OTUs revealed a marked change in community composition between the bioturbated samples and the subsurface below, producing a similar pattern as was seen for the total community (Figure 3c). The surface samples were dominated by OTUs classified as Uncultured Desulfobacteraceae lineage A (Jochum et al., 2017) and Syntrophobacteraceae-like DsrAB, which collectively comprised nearly 60% of the total sequences at 3 cmbsf (Figure 3d). While the relative abundances of these lineages decreased with depth, OTUs belonging to the Aarhus Bay lineage (Jochum et al., 2017) and the Uncultured Desulfobacteraceae lineage F (Jochum et al., 2017) became increasingly dominant downcore (Figure 3d). OTUs classified within the Aarhus Bay lineage comprised less than 0.01% of sequencing reads within the bioturbation zone, but increased to nearly 40% of the total sequencing reads at the bottom of the core (Figure 3d).

Absolute abundance estimates were calculated for each lineage by multiplying *dsrB* gene copy numbers by relative sequence abundances, assuming one copy of the gene per cell (Wagner et al., 2005). Figure 5 displays the three most prominent depth-associated trends in absolute abundances amongst the different lineages. Among these, we observed a marked drop in the abundance of OTUs belonging to the Uncultured Desulfobacteraceae lineage A, which declined by nearly one order of magnitude between the sediment surface and 7 cmbsf. This was a common trend for other predominant lineages, with substantial decreases in population size occurring for 8 out of the 15 most abundant SRM lineages (Supplementary, Figure S5). This steep drop in population sizes occurred in parallel to the rapid population growth of the Desulfobacteraceae lineage F, which took over as the dominant lineage below the bioturbation zone. This lineage was depleted again near the bottom of the SR zone (26-48 cmbsf), reaching a minimum abundance of 1.4 × 10^6^ (gene copies g^−1^ sediment). Within the same depth interval, members of the Aarhus Bay lineage became quantitatively dominant, reaching a maximum abundance of 2.1× 10^6^ (gene copies g^−1^ sediment) at 50 cmbsf. (Figure 5).

**Figure 5.**
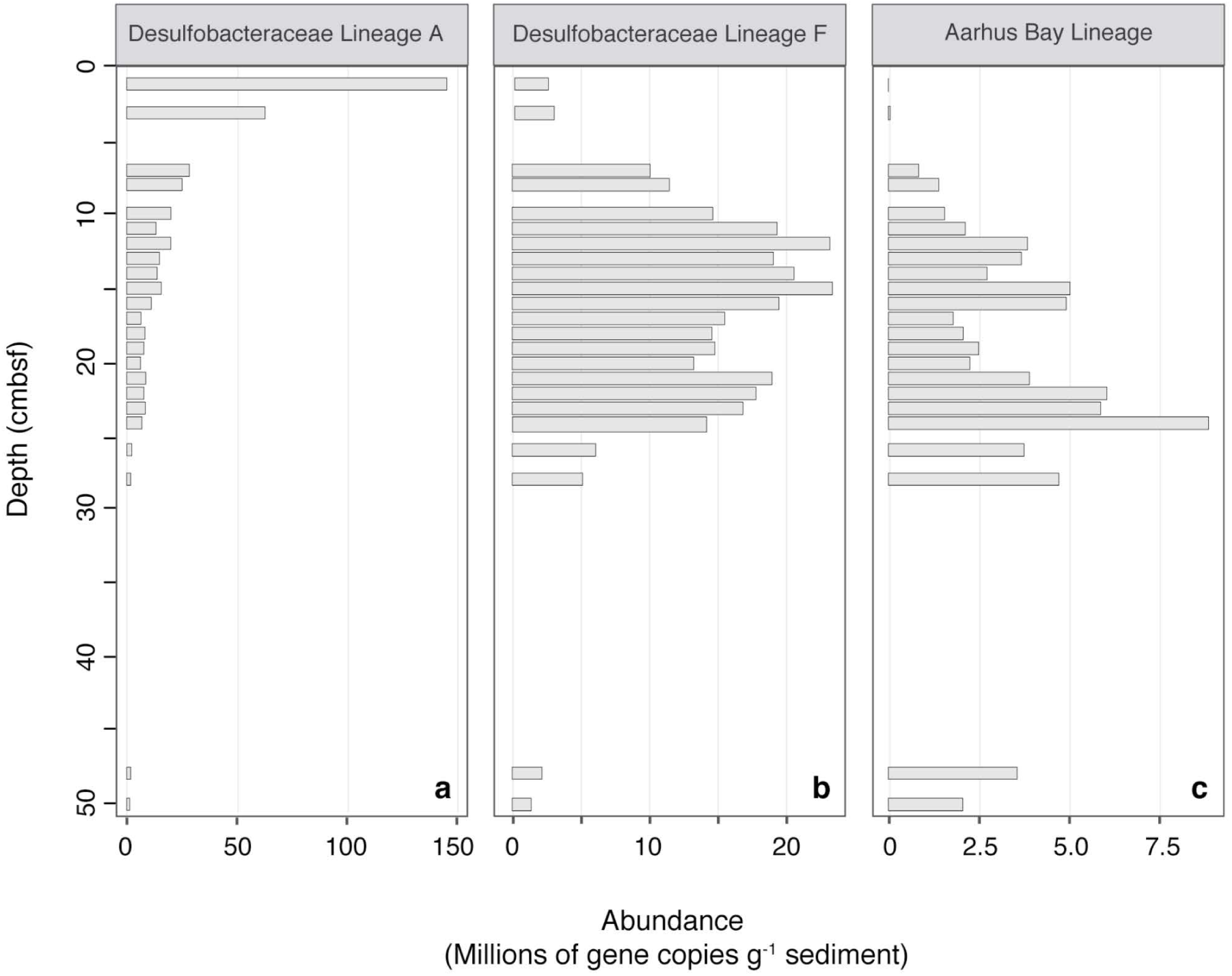
Absolute abundance profiles of SRM lineages with depth. Absolute abundances were estimated by multiplying sequence percent abundances by *dsrB* gene copy numbers, as determined by qPCR. Blank spaces indicate a lack of sequencing data.

### Cell-specific SRR and community turnover

Measurements of SRR and SRM abundance were used to estimate mean cell-specific metabolic rates (csSRR) throughout the core. The abundance of the total SRM community was estimated by dividing *dsrB* gene copy numbers by 7%, corresponding to the calculated DNA extraction efficiency. The csSRR were highest at the sediment surface, peaking at 0.07 fmol cell^−1^ day^−1^ at 3 cmbsf, and then decreasing towards a more constant value at 30-50 cmbsf (Figure 6a). This pattern was reproduced by samples taken across broader depth intervals in 2017 and 2018, indicating that microbial activity in the system is relatively stable over years (Figure S6).

**Figure 6.**
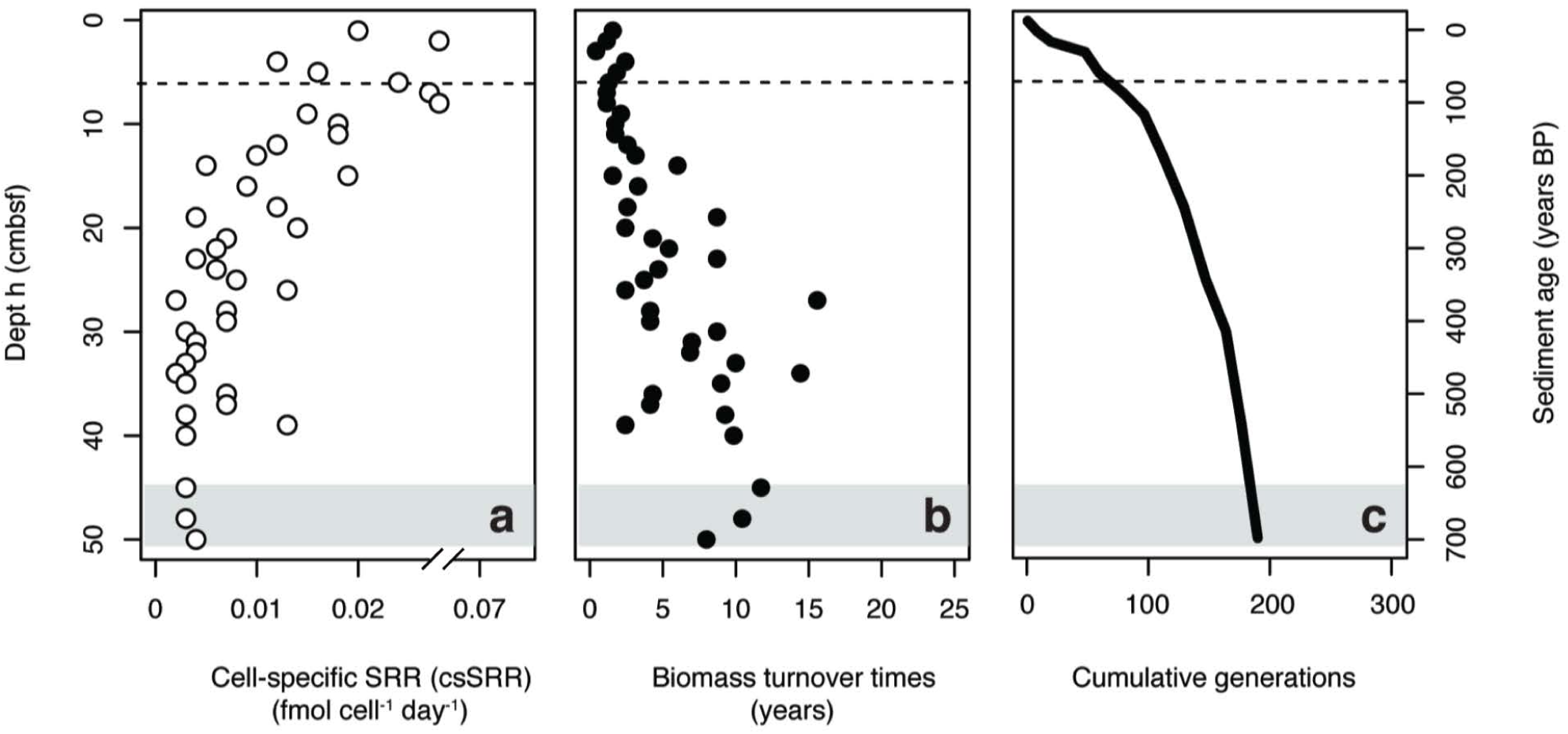
(a) Estimates of cell-specific sulfate reduction rates (csSRR) in the sediment core. (b) Estimated biomass turnover times of the total SRM community (c) Cumulative generations possible throughout the sediment core. All values were estimated using *dsrB* gene copy numbers as a proxy for SRM abundance, assuming a 7% DNA extraction efficiency. Gray shading indicates the onset of the SMT. Dashed lines in the surface indicate the bottom of the bioturbation zone.

Rates of SR were then used to calculate the amount of time required for complete turnover of SRM biomass (T_b_) within a given depth interval. Turnover times increased with depth below the bioturbation zone (Figure 6b), with an average value of 6 ± 4 years. Biomass turnover times were used to generate a profile of the number of cumulative generations that a community could undergo during burial (Figure 6c). The resulting profile indicated that a large fraction of possible generations within the depth profile could occur within the uppermost centimeters of the sediment column. Once buried below the bioturbation zone, the community would have undergone over 40% of the total generations possible within the 50 cm depth interval.

## 4. Discussion

### The influence of bioturbation on community assembly

Four processes—diversification, dispersal, selection, and drift, have been proposed as major drivers of microbial community assembly (Nemergut et al., 2013). These processes have been examined within the context of the marine subsurface environment, suggesting that subsurface microbial communities predominantly assemble by selective survival of buried taxa (Jochum et al., 2017; Petro et al., 2017; Starnawski et al., 2017). Genetic diversification is limited from the sediment surface down to 2 mbsf, suggesting that microbial communities do not undergo adaptive evolution during burial (Starnawski et al., 2017). This lack of apparent evolution in the seabed is expected to be due to exceedingly long generation times which increase with sediment depth and age (Hoehler and Jørgensen, 2013; Jørgensen and Marshall, 2016; Lever et al., 2015a; Røy et al., 2012). Our estimates of microbial activity and biomass turnover show that the highest rates of activity, and likewise the fastest biomass turnover times, occur within the most heavily bioturbated surface layer (1-6 cmbsf) of Aarhus Bay sediments. As communities are buried below the bioturbation zone, biomass turnover times increase steadily, limiting the number of possible generations and likewise the potential for genetic diversification (Figure 6). Due to the lack of diversification even over longer depth intervals (Starnawski et al., 2017), it would seem that the greatest scope for adaptive evolution in the seabed occurs within the uppermost 6 cm of the sediment column of Aarhus Bay. The occurrence of relatively high activities and community generations within the bioturbation zone can be explained by macrofaunal reworking, which introduces labile organic matter from overlying water and the seafloor into deeper sediment layers (Chen et al., 2017; Kristensen and Holmer, 2001; Kristensen and Kostka, 2013). Bioirrigation of sediment during burrowing, feeding, and respiration can further stimulate organic matter degradation by introducing high potential electron acceptors into the sediment (Aller and Aller, 1998).

Much like diversification, the influence of dispersal, or the movement of microbial cells in space, may be higher within the bioturbation zone than the subsurface below. The potential for passive dispersal was especially pronounced within the uppermost 6 cm of the sediment, due to homogenizing mixing by macrofauna (Figure 1). Below the bioturbation zone, mixing decreases by at least two orders of magnitude. When conditions conducive to passive transport largely cease and community turnover is diminished, selection caused by fitness differences between taxa (Nemergut et al., 2013) is likely to take over as the predominant means of assembly.

Our high depth resolution datasets including a mapping of the extent of bioturbation provides direct evidence that stark changes in microbial abundance and community structure occurs exactly at the bottom of the bioturbation zone (Fig. 1-3). As discussed in detail below, our results thus confirm the hypothesis that this is where the subsurface microbial community assembles in the sediment. This can be explained by a shift in the selection regime—from fast growth and adaptation to dynamic conditions (including exposure to O_2_) within the mixed surface, to survival under energetic limitations in the subsurface below. This change causes an increase in the relative abundances of taxa which have been found to predominate in much deeper sediments, where rates of biomass turnover may approach hundreds to thousands of years (Hoehler and Jørgensen, 2013; Jørgensen and Marshall, 2016; Parkes et al., 2000, 2014). In agreement, selection has been previously implicated as a major driver of microbial community assembly in the subsurface, first in aquifers (Stegen et al., 2013), and more recently in marine sediments (Jochum et al., 2017; Petro et al., 2017; Starnawski et al., 2017; Walsh et al., 2015). Here we see that selection for taxa adapted to energetic limitations begins already centimeters below the sediment surface, as the community becomes buried underneath the more dynamic and energy-rich bioturbation zone. While the most marked difference in community composition occurs between the bioturbated samples and the subsurface below (7 cmbsf), there is still variability in the composition of samples from 7-50 cmbsf (Figure 3). These differences suggest that the deep community is not yet fully assembled, and that gradual population changes are likely to continue to occur throughout the entirety of the sediment column.

### Influence of geochemical zonation

The distributions of abundant SRM lineages were indicative of a response to geochemical zonation, with major shifts in abundances occurring at the onset and end of the SR zone (Figure 5). The lineages that increased in absolute abundance within the SR zone maintained relatively constant abundances throughout the entirety of this region, with gene copy numbers dropping upon transition into the SMT. Apparent responses to geochemical zonation were especially pronounced for Uncultured Desulfobacteraceae lineage F and the Aarhus Bay lineage—both of which have been found previously to dominate the SRM community below the sediment surface and down into the methanogenesis zone (Jochum *et al.*, 2017). The vertical stratification of SRM taxa has been demonstrated previously in coastal sediments (Sahm et al., 1999), biofilms (Ramsing et al., 1993), and marine Arctic sediments (Ravenschlag et al., 2000). These observations can be explained by an adaptation of certain lineages to specific geochemical conditions, allowing them to become dominant within a given depth interval (Jorgensen et al., 2012). The assumption that community dynamics are reflective of different ecological niches among the lineages is further supported by the measured SRR, which reach maximum rates at 3 cmbsf (Figure 1b). This delay, which is indicative of the competitive inhibition of dissimilatory sulfate reduction by Mn and Fe reductive processes (Canfield et al., 1993; Thamdrup et al., 1994), suggests that sulfate reduction is not the predominant terminal degradation process in the top 1-2 cm of the sediment. The geochemical succession of specialized populations is nicely illustrated by the *dsrB* sequence dataset, which shows that the dying off of dominant taxa within the uppermost 1-3 cm depth interval is followed by the rapid growth of lineages that subsequently take over as predominant members of the community (Figure 5).

While geochemical zonation thus has a hand in regulating some changes within the SRM community between depths, the overall trends for both SRM and total community appear decoupled from the geochemical zonation (Figure 3). This observation implies that the availability of energy and carbon, and not the concentration of electron acceptors such as sulfate, is the major driver for the microbial community, including also the terminal oxidizers.

### Persisting sediment community

Extensive molecular surveys of marine sediments have identified a unique assemblage of microorganisms, which comprise a significant portion of the deep subsurface community at geographically distinct locations (Fry *et al*., 2008; Webster *et al*., 2004; Teske and Sørensen, 2008; Inagaki *et al*., 2003; Orcutt *et al*., 2011; Walsh *et al*., 2015). Many of these dominant taxa have been found to belong to a subset of ‘persister’ microorganisms, whose relative abundances increase with sediment depth, irrespective of geochemical zonation (Petro et al., 2017; Starnawski et al., 2017). Common persisting lineages within the Bacteria include the Atribacteria (Dodsworth et al., 2013; Nobu et al., 2015) or Chloroflexi, while the archaeal community is dominated by members of the Bathyarchaeota (Meng et al., 2014) and Lokiarchaeota (Inagaki et al., 2003; Spang et al., 2015; Vetriani et al., 1999). While these taxa are commonly associated with deep marine sediments, here we see that their increase in relative abundance begins just centimeters below the seafloor, right under the bioturbation zone (Figure 3). The persistence of OTUs throughout the depth profile was demonstrated by tracing OTUs across depths in the sediment (Supplementary Figure S2). A small subset (<100) of OTUs were found to persist from the surface down to 50 cmbsf, where they came to comprise >40% of the total community. Similar patterns of OTU overlap have been observed between sediments and the overlying seawater, suggesting that deep subsurface communities are comprised of rare taxa that are deposited from the water column and persist throughout burial (Walsh et al., 2015). Our estimates of the absolute abundances of dominant OTUs demonstrate that select populations may increase in absolute abundance with sediment depth (Figure 4), contrary to the drop in cell numbers seen for the total community. This suggests that persisting populations may not simply survive, but also grow, within the energy limited subsurface. While striking, this observation is based on the relative abundances of OTU sequences combined with qPCR data of the total population, and should be verified using direct quantitative methods, such as fluorescence *in situ* hybridization (FISH) or OTU-specific qPCR, in future studies.

While the data presented here are only collected from a single sampling site, similar patterns of microbial diversity have been observed across broader sediment depths at numerous sites within Aarhus Bay (Jochum et al., 2017; Starnawski et al., 2017). By finely resolving population dynamics near the sediment surface at site M5, we see that a marked shift in the composition of the subsurface community occurs already centimeters below the seafloor, immediately below the bioturbation zone. The taxa which persist to the bottom of the sediment core are also found in low relative abundances at the surface, suggesting that populations present within the bioturbation zone act as a seed community for the subsurface below. Replicate cores sampled in 2017 and 2018 demonstrate that these depth-associated changes in the community are consistent across time. The geochemistry and rates of sulfate reduction were also stable across sampling dates, indicating a consistent drop in microbial activity between the bioturbation zone and the unmixed sediment below.

Collectively, these results suggest that the microbial communities present within the deep biosphere in Aarhus Bay sediments begin to assemble below the bottom of the bioturbation zone, where sediment mixing and energy availability are both diminished. These changes delineate the bioturbation zone from the unmixed sediment below, where environmental selection for populations adapted to energy limitations starts to shape the microbial communities which will come to predominate within the energy-starved deep subsurface biosphere in hundreds to thousands of years.

## 5. Acknowledgements

We thank the crew and scientists on board of RV Aurora for help with sampling as well as Britta Poulsen, Susanne Nielsen, and Jeanette Pedersen for significant technical assistance in the laboratory. We thank Ian Marshall for providing us with a Python script used for our sequence analyses, and for helping, together with the participants of the course ‘Microbial Element Cycling and Population Ecology’, to retrieve microbial community data in 2017 and 2018.

## 7. Author Contributions

BJ, HR, KK, and AS designed the research. CP, BZ, and TF performed the research. CP, BZ, LJ, PS, and HR analyzed the data. CP and HR wrote the paper with input from all coauthors.

