## Supplementary Material for "Marine deep biosphere microbial communities assemble in near-surface sediments in Aarhus Bay"

***Analysis of sediment cores from 2017 & 2018***

In order to assess the reproducibility of the trends observed in our fine-scale analysis of a single core, four replicate sediment cores were taken from site M5 in Aarhus Bay during both 2017 and 2018. These cores were analyzed with a broader depth resolution than the core from 2014, focusing only on the sediment surface (1-2 cmbsf), the bottom of the bioturbation zone (5 cmbsf), and below (10-50 cmbsf) in 10 cm depth intervals. Within each sample, we analyzed (1) microbial activity (SRR), (2) microbial abundances (qPCR), and (3) community composition (Illumina MiSeq sequencing).

SRR and qPCR were both performed as described in the primary text, while sequencing of the 16S rRNA gene and *dsrB* gene were performed with a modified protocol and a different sequencing techonology. Bacteria and Archaea were sequenced separately, using the primer pairs Bac341F/Bac805R (Herlemann et al., 2011) and Arch344Fmod/Arch915R (Casamayor et al., 2002; Xiao et al., 2017), respectively. To conserve time and resources, Bacteria and Archaea were sequenced separately in 2017 (Bacteria) and 2018 (Archaea). Amplicon libraries for Bacteria and Archaea were preprared and sequenced according to Vergeynst *et al.* (2018) and Xiao *et al.* (2017), respectively. All libraries were sequenced on an Illumina MiSeq system at Aarhus University (Aarhus, Denmark). The raw sequencing reads were processed using the package DADA2 (v1.5.0) and the standard DADA2 workflow (Callahan et al., 2016) in R. Amplicon sequence variants (ASVs) were taxonomically classified in DADA2 using the SILVA SSU Ref v1.32 (Quast et al., 2013).

| A | | | | | |
|---|---|---|---|---|---|
| **OTU** | **Phylum** | **Class** | **Order** | **Family** | **Genus** |
| **8** | Proteobacteria | Gammaproteobacteria | Chromatiales | Ectothiorhodospiraceae | *Acidiferrobacter* |
| **9** | Proteobacteria | Deltaproteobacteria | Myxococcales | Sandaracinaceae | unclassified |
| **36** | Proteobacteria | Deltaproteobacteria | Desulfobacterales | Desulfobulbaceae | *Desulfobulbus* |
| **11** | Bacteroidetes | BD2-2 | unclassified | unclassified | unclassified |
| **100** | Proteobacteria | Deltaproteobacteria | Desulfobacterales | Desulfobacteraceae | Sva0081 sediment group |
| **39** | Proteobacteria | Deltaproteobacteria | Desulfuromonadales | Sva1033 | unclassified |
| **82** | Proteobacteria | Gammaproteobacteria | Alteromonadales | Alteromonadaceae | OM60(NOR5) clade |
| **14** | Proteobacteria | Gammaproteobacteria | Chromatiales | Ectothiorhodospiraceae | *Acidiferrobacter* |
| **18** | Gemmatimonadet es | Gemmatimonadetes | PAUC43f_marine benthic_group | unclassified | unclassified |
| **19** | Actinobacteria | Acidimicrobiia | Acidimicrobiales | OM1_clade | unclassified |
| **B** | | | | | |
| **OTU** | **Phylum** | **Class** | **Order** | **Family** | **Genus** |
| **10** | Atribacteria | unclassified | unclassified | unclassified | unclassified |
| **2** | Proteobacteria | Deltaproteobacteria | Desulfarculales | Desulfarculaceae | unclassified |
| **41** | Actinobacteria | OPB41 | unclassified | unclassified | unclassified |
| **43** | Planctomycetes | Phycisphaerae | MSBL9 | unclassified | unclassified |
| **12** | Deinococcus-Thermus | Deinococci | Thermales | Thermaceae | *Oceanithermus* |
| **73** | Proteobacteria | Deltaproteobacteria | Desulfarculales | Desulfarculaceae | unclassified |
| **47** | Other | unclassified | unclassified | unclassified | unclassified |
| **132** | Nitrospirae | Nitrospira | Nitrospirales | Nitrospirales_Incertae Sedis | *Candidatus* Methylomirabilis |
| **20** | Spirochaetae | Spirochaetes | Spirochaetales | Spirochaetaceae | Termite Treponema cluster |
| **1** | Candidate_divisio n_OP8 | unclassified | unclassified | unclassified | unclassified |

**Table S1.** Taxonomic identity of OTUs displayed in Figure 4. (a) OTUs that decrease in absolute abundance with sediment depth, displayed in Figure 4a. (b) OTUs that increase in absolute abundance with sediment depth, displayed in Figure 4b.

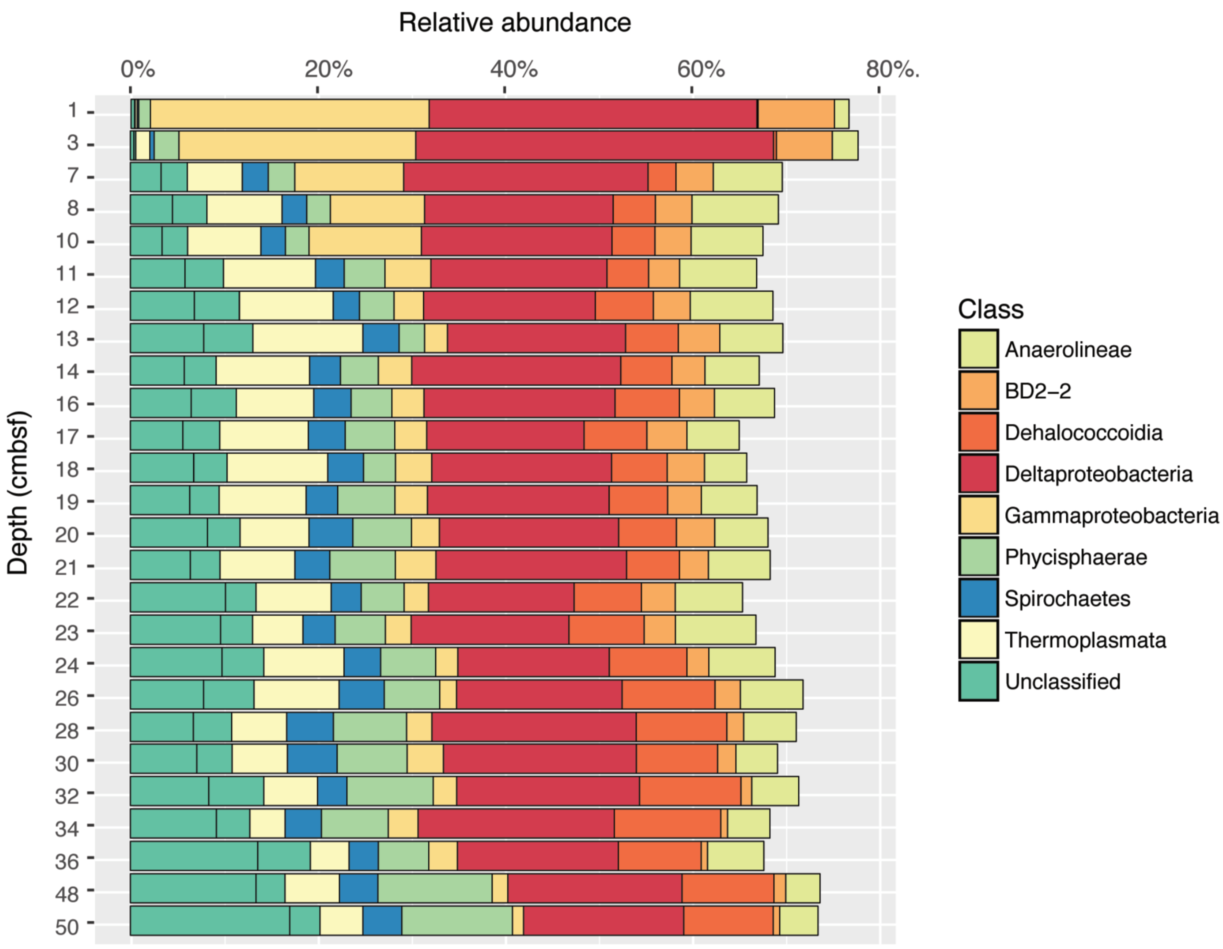


**Figure S1.** Relative abundance of the top 10 most abundant microbial classes identified by 16S rRNA gene sequencing. Colors correspond to the taxonomic identity at the class level and bar lengths represent the relative abundances of the classes out of the total number of sequencing reads at each depth.

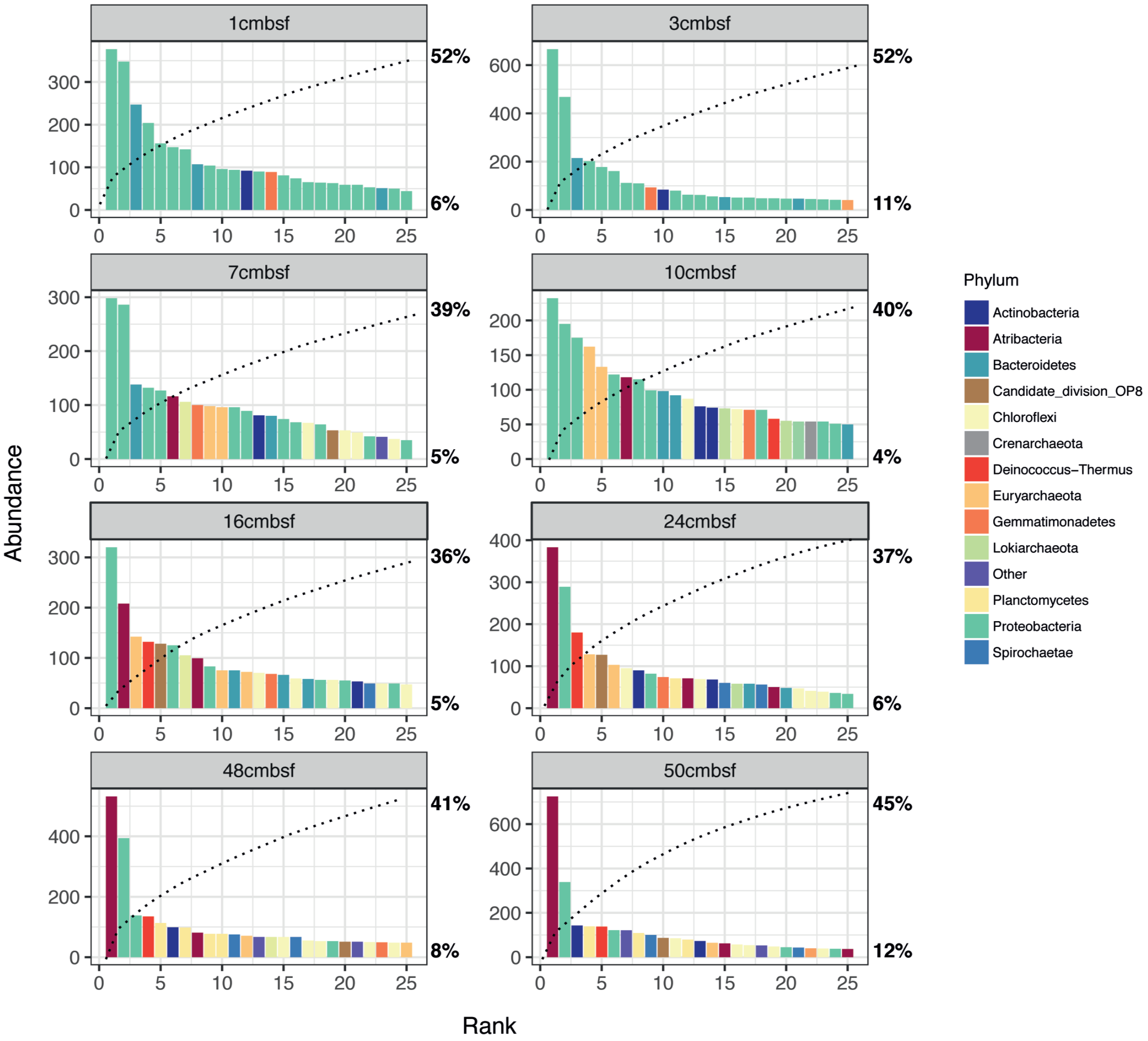


**Figure S2.** Rank abundance curves of the 25 most abundant OTUs identified by 16S rRNA gene sequencing throughout the depth profile. Each panel displays the 25 most abundant OTUs (97% identity cutoff) at the indicated depth. The x-axis shows the rank of each OTU and the y-axis shows the number of sequencing reads assigned to each OTU. The colors of the bars indicate the taxonomic identity of each OUT on the phylum level. The dotted lines show the cumulative relative abundance for the OTUs out of the total number of sequencing reads at that depth.

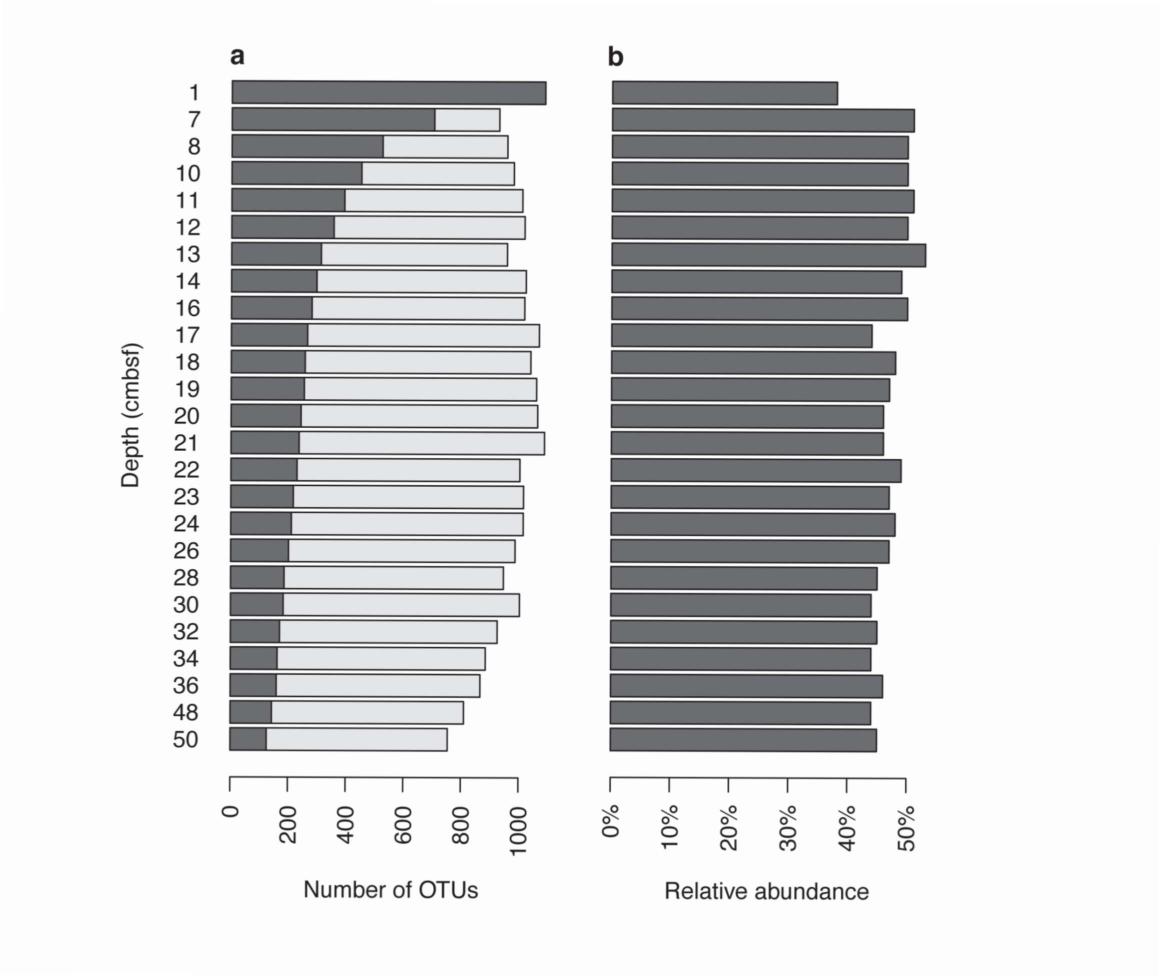


**Figure S3.** OTUs persisting across depths. (a) Abundance of 16S rRNA gene sequence OTUs that persist across all depths in the sediment core. Light bars show the total number of OTUs present within a depth interval. Dark grey bars show the number of OTUs present within a depth that were also present in all of the above depths. (b) Relative abundance of persisting OTUs at each depth.

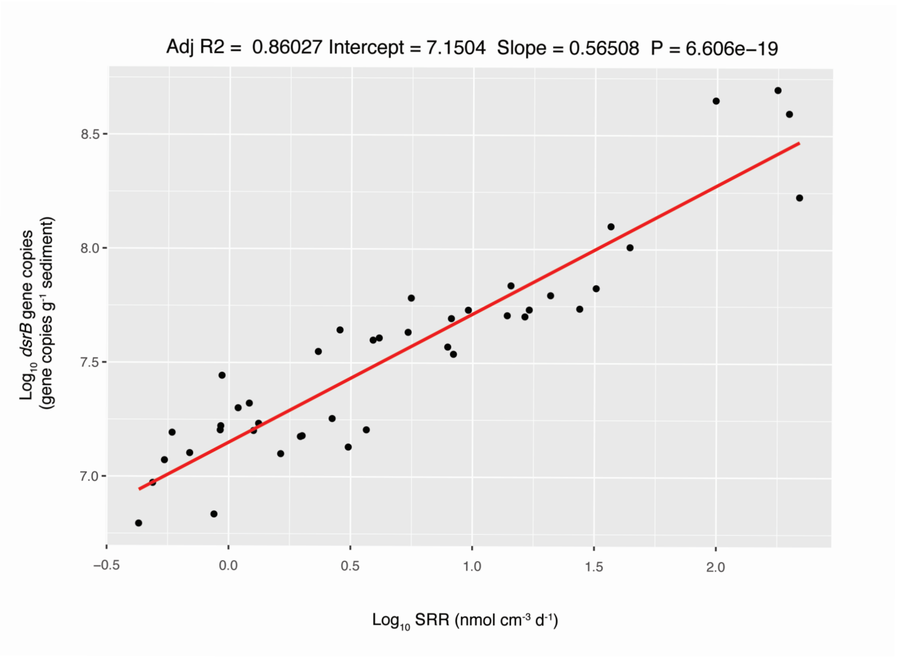


**Figure S4.** Correlation of dsrB gene copy numbers to measured sulfate reduction rates (SRR). P value is Pearson correlation coefficient.

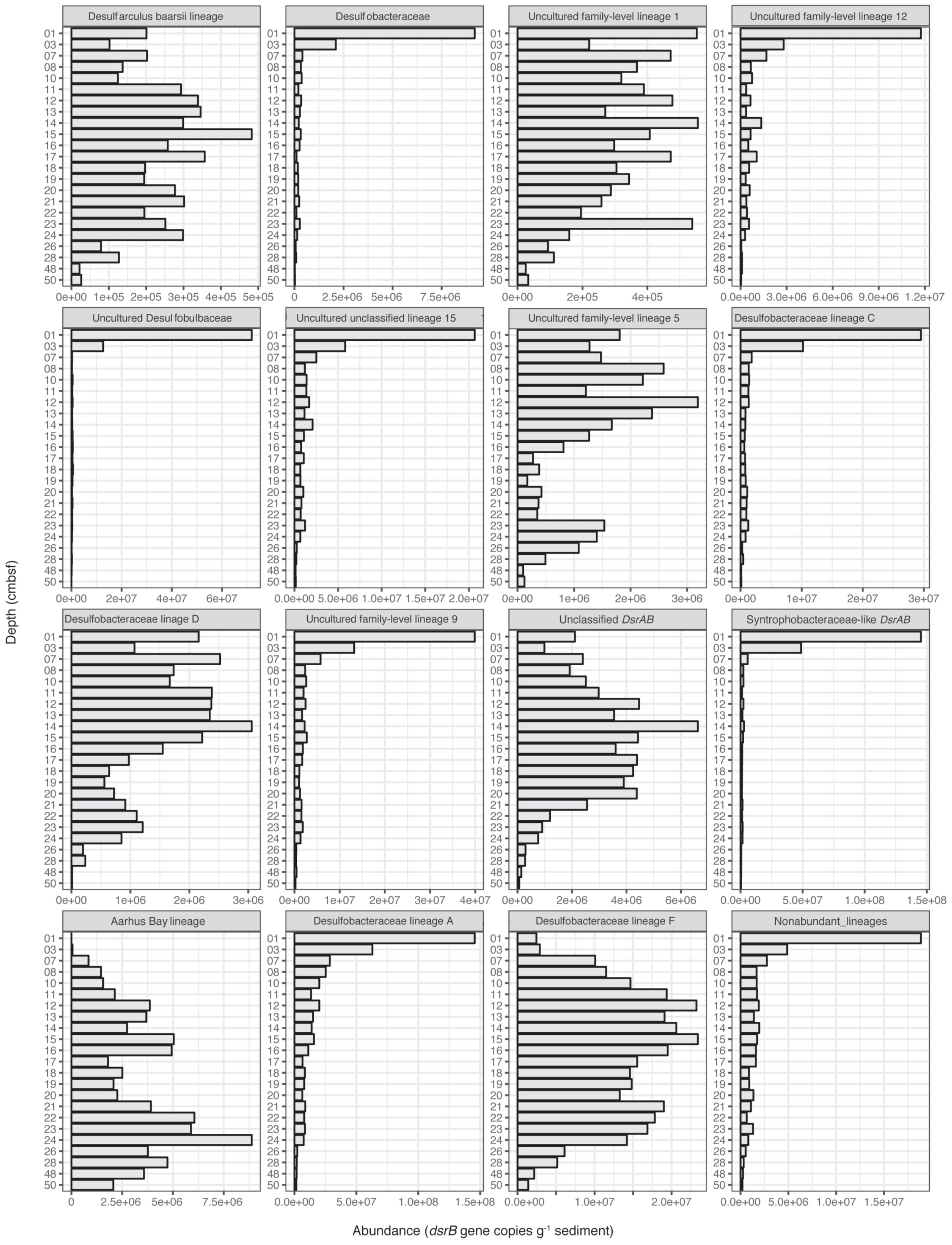


**Figure S5.** Absolute abundances of dominant lineages of sulfate reducing microorganisms (SRM). Absolute abundances were estimated by multiplying the relative abundances of dsrB gene sequences classified within each lineage by total dsrB gene copy numbers obtained from qPCR. Pooled nonabundant lineages comprised less than 20% of total sequences at each depth.

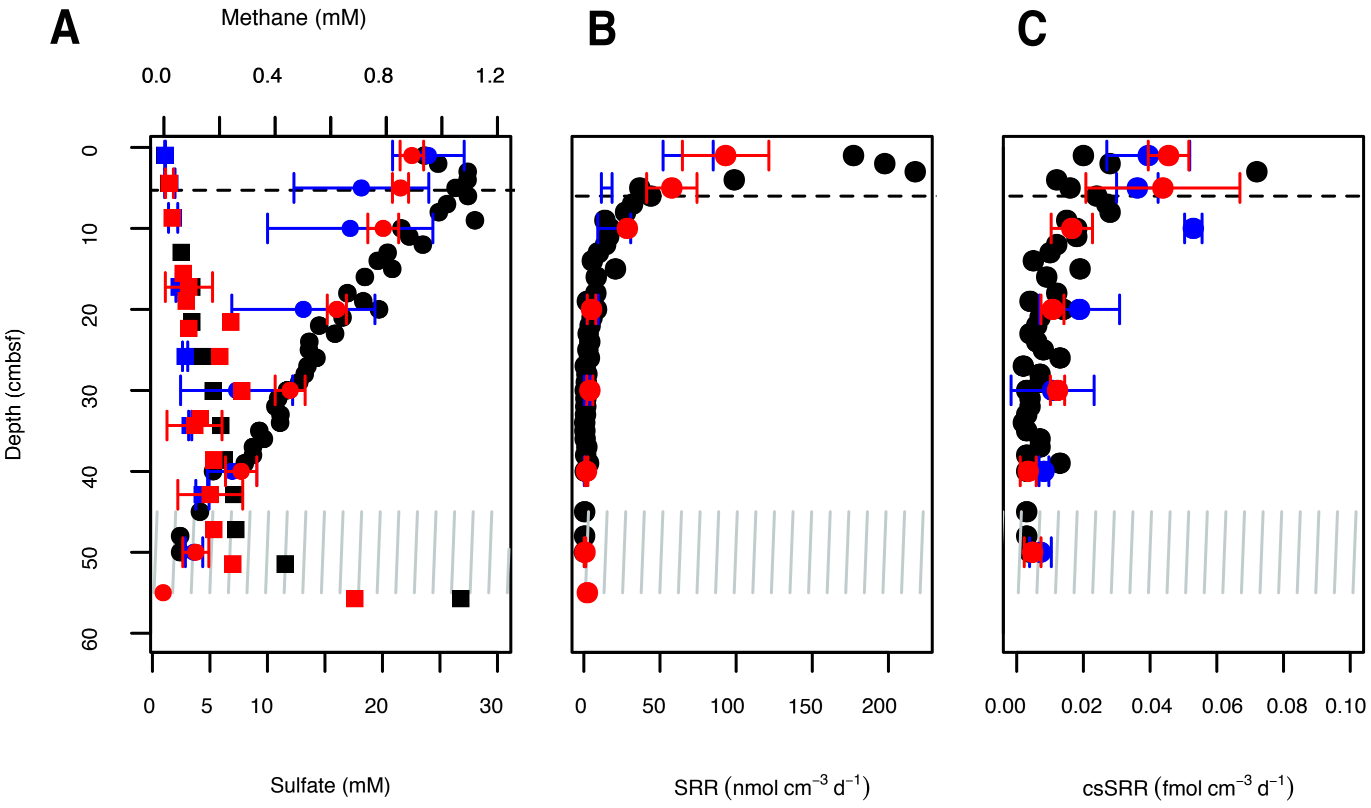


**Figure S6.** Spatiotemporal variation in (A) Sediment porewater concentrations of sulfate and methane, (B) sulfate reduction rates (SRR), and (C) cell-specific sulfate reduction rates estimated using dsrB gene copy numbers as a proxy for SRM abundance. DsrB gene copy numbers are displayed in Figure S8. For (A) sulfate data are shown by circles and methane data are shown by squares.

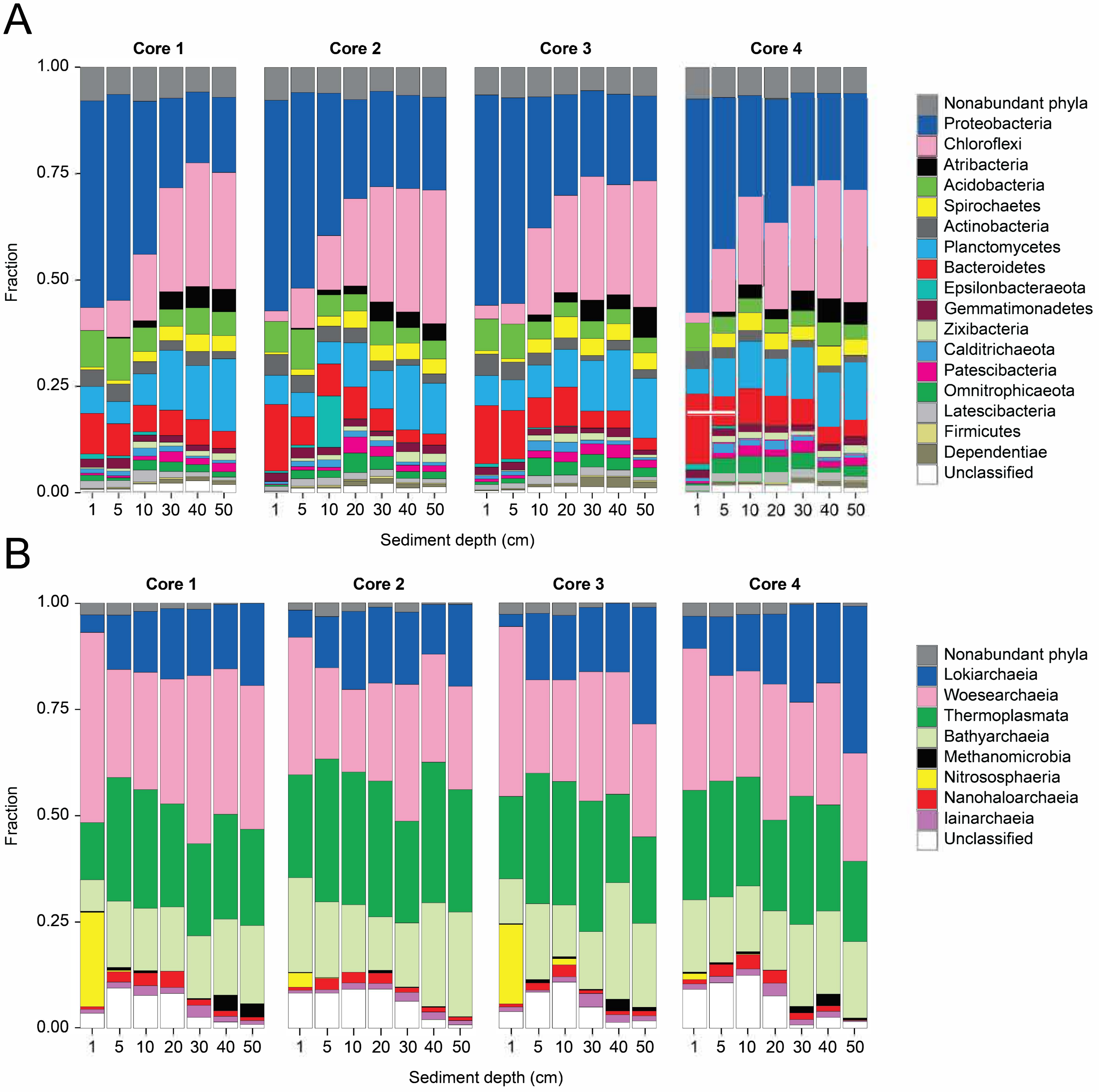


**Figure S7.** Microbial community composition within cores sampled in 2017 and 2018. The stacked bar plots show relative abundances of taxonomic groups identified by 16S rRNA gene sequencing. (A) Relative abundance of Bacteria on the phylum level from four replicate cores collected in 2017. (B) Relative abundance of Archaea on the class level from cores collected in 2018.

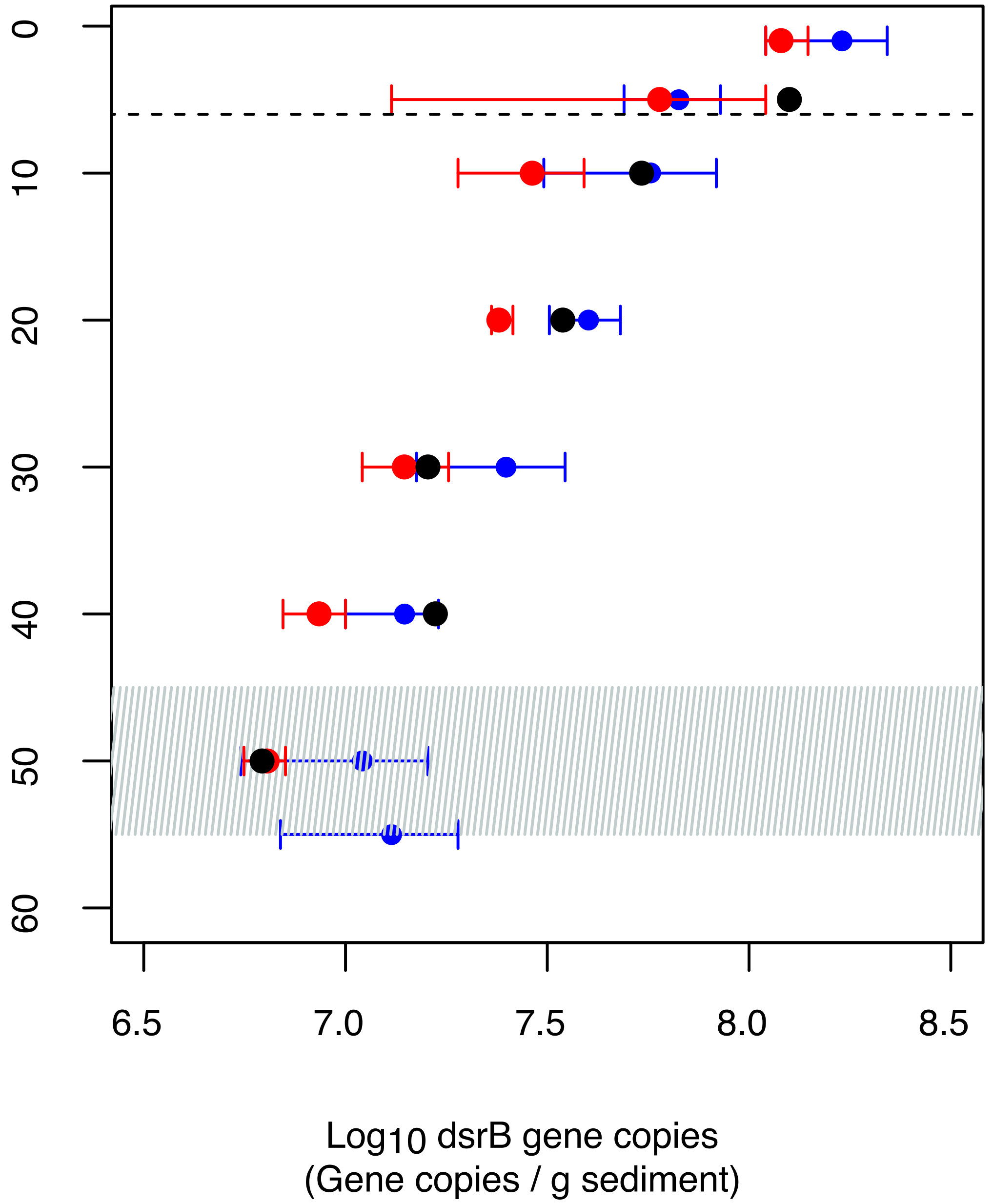


**Figure S8.** Spatiotemporal variation in abundance of SRM, as measured by qPCR of the dsrB gene. The color of the symbols refers to the sampling date, with cores taken from 2014 (black, n=1), 2017 (blue, n=4), and 2018 (red, n=4). The number of dsrB gene copies are presented on a log10 scale.
